# Functional single-cell genomics of human cytomegalovirus infection

**DOI:** 10.1101/775080

**Authors:** Marco Y. Hein, Jonathan S. Weissman

## Abstract

The complex life cycle of herpesviruses is orchestrated by the interplay of host factors and hundreds of viral genes. Understanding how they work together and how perturbations of viral and host factors impact infection represents both a fundamental problem in virology and the basis for designing antiviral interventions. Here, we use CRISPR screening to comprehensively define the functional contribution of each viral and host factor to human cytomegalovirus (HCMV) infection in primary cells. We then record the transcriptomes of tens of thousands of single cells, and monitor how genetic perturbation of critical host and viral factors alters the timing, course, and progression of infection. We find that normally, the large majority of cells follow a stereotypical transcriptional trajectory. Perturbing critical host factors does not change this trajectory per se, but can either stall, delay or accelerate progression along the trajectory, allowing us to pinpoint systematically the stage of infection at which each host factor acts. Conversely, perturbation of viral factors can create distinct, abortive trajectories. Our results reveal a dichotomy between the roles of host and viral factors and more generally provide a road map for functional dissection of host-pathogen interactions.

---

The β-herpesvirus HCMV is a pervasive pathogen that establishes lifelong infection in the majority of the human population. Activation of its lytic cycle triggers a characteristic cascade of events, starting with stereotypical waves of viral gene expression, continuing with the replication of the large, 235 kb dsDNA genome, and culminating in the budding of newly assembled virions (1). A number of systems-level studies have described these phenomena on the level of the transcriptome, the set of translated messages, and the proteome in time and space (2–7). A core set of viral genes essential for replication has been established by systematic mutagenesis of the viral genome (8, 9). These studies have highlighted the complexity of the viral replication cycle and have raised the question of how hundreds of viral genes cooperate to manipulate the host and undermine its defense machinery, and which are the most promising targets for antiviral intervention.

CRISPR-Cas9 technology provides us with tools to systematically measure the functional contribution of each viral gene and host factor involved in productive infections (10). However, it remains a challenge to translate a list of candidate factors identified in a screen into a systematic understanding of the roles that individual factors play during viral infection and how they are organized in distinct pathways. We address this challenge with a scalable approach that combines CRISPR-based genetic perturbations with rich phenotypic profiling by single-cell transcriptomics. First, we conducted systematic pooled CRISPR screens for both host and viral factors affecting survival of primary human fibroblasts upon HCMV infection. A simple readout such as cell survival makes pooled screens scalable to genome-wide libraries, but captures only a compressed picture of the complexity of molecular events impacted by altering expression of host or viral factors. Moreover, lytic infection is inherently dynamic over time, and heterogeneous from cell to cell in a population (11, 12). We therefore recorded the transcriptomes of tens of thousands of single cells during infection, and monitored how perturbation of a set of critical host and viral factors—identified in the pooled screens—alters the timing, course, and progression of infection at single-cell resolution. Our data paint a high-resolution picture of the HCMV lytic cycle as a deterministic program that is distinctly vulnerable to host- and virus-directed interventions: We identify host dependency factors critical for viral entry and the progression from early to late stage of infection, as well as restriction factors that slow the pace of progression. Conversely, we show that targeting key viral factors derails the viral gene expression program in specific ways. Taken together, our findings reveal a dichotomy between the roles of host and viral factors, with the set of viral factors solely defining the trajectory of infection and host factors creating the environment permitting the execution of that program.

## High resolution functional scanning of the HCMV genome reveals an architecture of functional modules

Several observations argue that Cas9 represents an effective tool for making targeted disruptions in the cytomegalovirus genome. Targeting individual essential herpesvirus genes by CRISPR/Cas9 was shown to disrupt their expression directly and their function through errors introduced by the host DNA repair machinery (13). Cleavage of the viral DNA in non-essential regions has minimal impact on HCMV replication and host cell viability—likely because DNA repair is fast relative to the kinetics of replication—but can affect expression of genes proximal to the cut sites ((13) and our data below). To enable high-resolution scanning of viral elements for a comprehensive functional annotation of the HCMV genome, we designed a single-guide RNA (sgRNA) library that targets every protospacer-adjacent motif (PAM) for *S. pyogenes* Cas9 (NGG PAM sequence present roughly every 8 bp) along the genome of the clinical HCMV strain Merlin (Figure 1A, Table S1, Methods). We delivered the library into primary human foreskin fibroblasts engineered to express Cas9, so that upon HCMV infection, each cell executes a cut at a defined position along the viral genome, collectively tiling its entirety.

**Figure 1.**
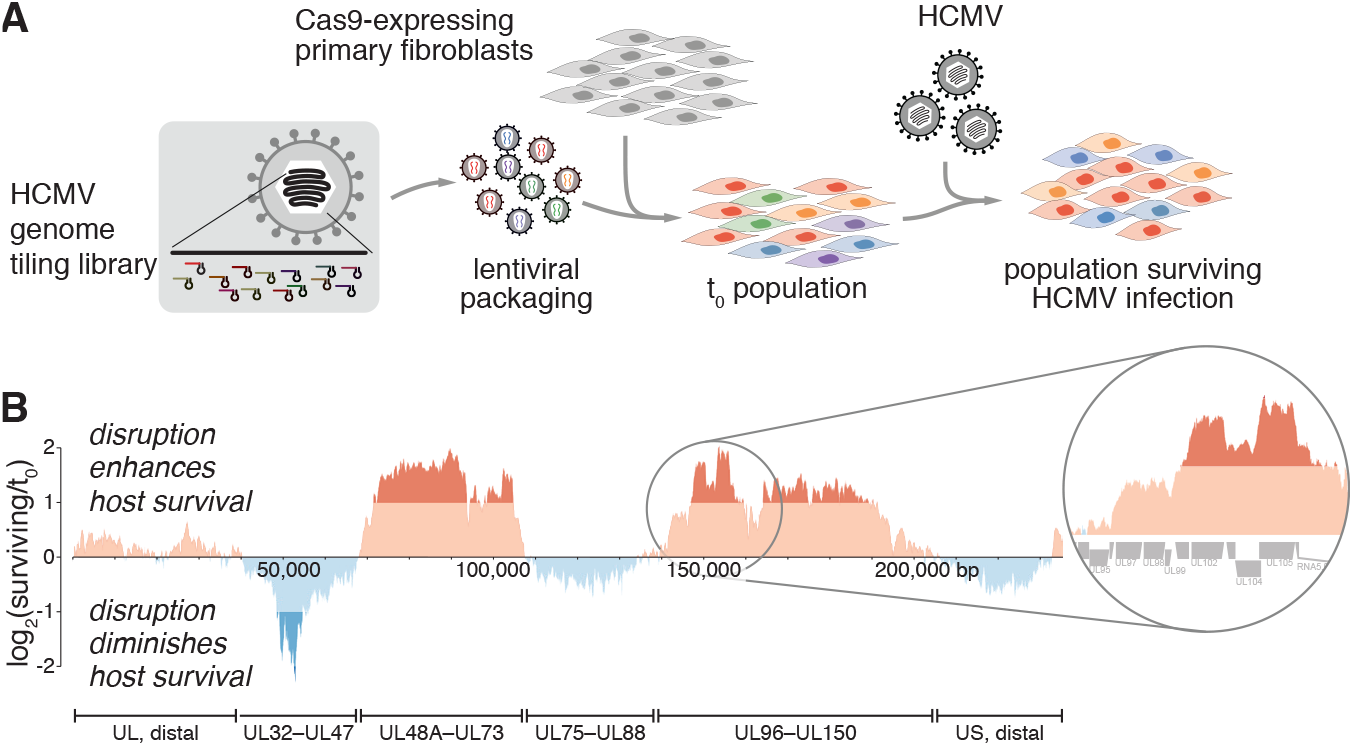
Virus-directed CRISPR nuclease screen maps the phenotypic landscape of the HCMV genome. (**A**) Experimental design for pooled, virus-directed CRISPR screening. Our HCMV tiling library contains ∼33k elements and was lentivirally delivered into primary human foreskin fibroblasts expressing the CRISPRn machinery, followed by infection with HCMV. sgRNA cassettes were quantified by deep sequencing in the initial (t_0_) population and the surviving population. (**B**) Phenotypic landscape of the HCMV genome obtained by locally averaging the phenotypes of individual sgRNAs. Strong changes in the magnitude of the phenotype coincide with gene-gene boundaries (inset).

We mapped the functional landscape of the HCMV genome by quantifying the abundances of individual sgRNA cassettes in cells surviving infection relative to the initial population by next-generation sequencing (Figure S1). We found that cutting phenotypes are relatively constant within individual genes, i.e. the determining factor is which gene is targeted by Cas9, more so than the relative position of the target site within the gene body. Cutting adjacent sets of genes frequently showed similar phenotypes. However, some gene boundaries were marked by abrupt phenotype changes, arguing that strong functional consequences of Cas9-induced double-strand breaks are limited to the vicinity of the cut sites (Figures 1B, S1, Table S1).

At a larger scale, changes in the direction and magnitude of the phenotypes defined six major genomic modules: Cuts in both distal regions of the genome, which lack genes essential for viral replication (8, 9), had minimal impact on host cell survival. As expected, targeting the two regions covering UL48A–UL73 and UL96–UL150, both of which contain essential genes involved in viral DNA replication, packaging and nuclear egress (9, 14), strongly protected infected cells from death. In the two remaining regions of the genome, we found that disruption of genes required for viral replication did not necessarily protect the host from death. Cuts within the UL32–UL47 region, which contains genes essential for viral replication, actually led to a strongly increased ability of the virus to kill cells. The most strongly sensitizing phenotypes mapped to the known viral apoptosis inhibitors UL36, UL37, and UL38 (15). While this behavior can be rationalized for virally encoded anti-apoptotic proteins, it extended to many other virus-essential genes without known anti-apoptotic roles, including the DNA polymerase processivity factor UL44. Finally, and counterintuitively, cuts in the central region spanning UL75–UL88 caused very mild phenotypes. Many genes in this region encode essential structural components of the viral envelope, tegument, and capsid, yet the outcomes on host survival after Cas9 cutting are comparable to targeting nonessential genes in the US distal region, resulting in even mildly enhanced cell death upon infection.

Targeting essential viral genes, by definition, undermines the production of viral offspring. The outcome for the host, however, is more nuanced and sometimes counterintuitive. It appears that disrupting essential genes involved in viral DNA replication mostly protects the host. However, interfering with the later steps of assembling new virions may not only be ineffective for protecting the host, but even place an additional burden.

## Genome-wide screens for host factors of HCMV infection define pathways affecting host survival

Next, we carried out a pooled screen for host factors modulating HCMV infection by systematically repressing expression of human genes by CRISPR interference (CRISPRi) (16, 17). Phenotypes were defined by enrichment or depletion of sgRNA cassettes in the surviving cell population over the initial population, as well as in an uninfected control population to account for the impact of a gene’s essentiality (Figure 2A, Table S2).

**Figure 2.**
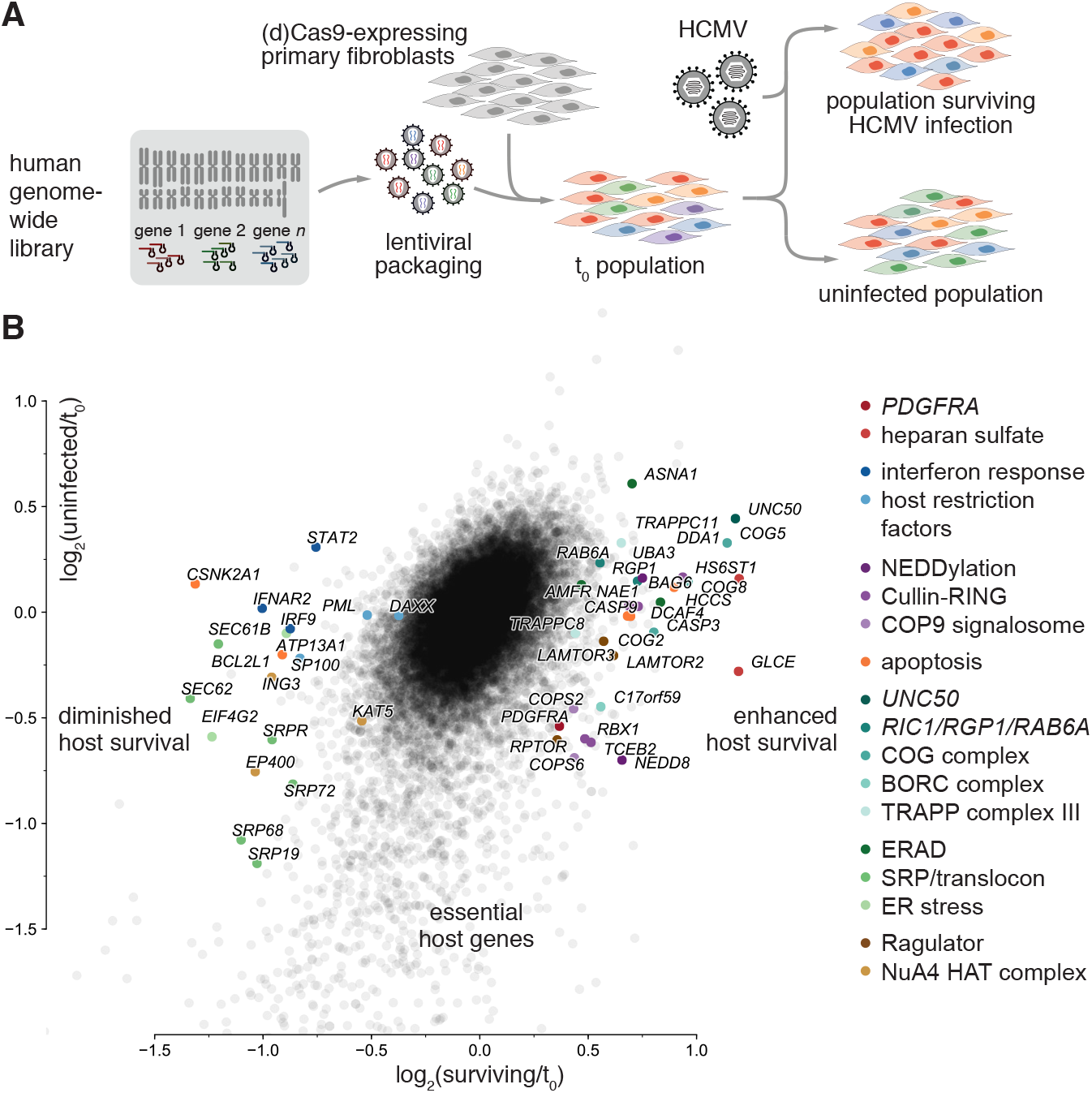
Host-directed CRISPR screens identify host dependency and restriction factors. (**A**) Experimental design for pooled, host-directed CRISPRi/n screening. Genome-wide sgRNA libraries targeting human genes with multiple sgRNAs each were lentivirally delivered into primary human foreskin fibroblasts expressing the CRISPRi/n machineries, followed by infection with HCMV. sgRNA cassettes were quantified by deep sequencing in the initial (t_0_) population, the surviving population and an uninfected control population to account for gene essentiality in absence of infection. (**B**) Results of the host-directed CRISPRi screen displayed as a scatter plot of average gene essentiality (i.e. infection-independent phenotype; y-axis) vs. protection/sensitization to death upon HCMV infection (i.e. infection-dependent phenotype; x-axis).

Our screen revealed a range of diverse host genes required for multiple steps in the viral life cycle (Figure 2B). Genes involved in the biosynthesis of heparan sulfate were among the strongest protective hits. Heparan sulfate proteoglycans on the cell surface enable the adhesion of HCMV (18, 19). Additionally, we found a range of vesicle trafficking factors: *RAB6A* and its GEFs *RIC1*/*KIAA1432* and *RGP1*, the conserved oligomeric Golgi (COG) complex, members of TRAPP complex III, and *UNC50*. These factors converge on the Golgi apparatus and mediate both retrograde and anterograde transport. Some of these factors (e.g. COG complex, TRAPP complex, *UNC50*) had previously been implicated in the internalization of diverse bacterial and plant toxins, suggesting that HCMV and toxins exploit similar pathways for cell entry (16, 20–24).

Other protective hits included members of the LAMTOR/Ragulator complex, Folliculin (*FLCN*), and the Lyspersin (*C17orf59*/*BORCS6*) subunit of the BORC complex, all linked to lysosome positioning and nutrient sensing (25–27). This supports and extends the recent observation that HCMV infection changes lysosome dynamics (7). Additionally, host cell death was reduced by knock-down of certain Cullin-RING E3 ligases (*RBX1, CUL3*), their adaptor subunits (*DDA1, TCEB2*/*ELOB*), substrate receptors (*DCAF4*), and the associated neddylation (*NEDD8, NAE1, RBX1*) and deneddylation machineries (COP9 signalosome complex members). Many viruses, including HCMV, hijack this pathway to degrade host restriction factors, which can be prevented by broadly-acting Nedd8-activating Enzyme inhibitors (28, 29). Finally, we identified genes involved in tail-anchored protein insertion into the ER, as well as ER-associated degradation: *AMFR*, an E3 ligase, and the TRC40/GET pathway members *BAG6* and *ASNA1*, which were shown to be required for insertion of membrane proteins of herpes simplex virus 1 (HSV-1) which, however, lack HCMV orthologs (30).

Our screens also identified a number of host restriction factors, whose knockdown sensitizes cells to death upon infection rather than protecting them. Among these were known factors such as *PML* and *DAXX*, as well as members of the interferon type I (IFN) pathway. Sensitizing hits also included subunits of the NuA4 histone acetyltransferase complex, which was shown to counteract Hepatitis-B virus replication by binding to chromatinized viral DNA, acting as a transcriptional repressor (31). It has also been described as interactor of the HIV-1 TAT protein (transactivator of transcription) (32), but has not been described as herpesviral host factor. Further, we identified members of the signal recognition particle, the translocon and associated factors, and genes involved in ER stress (33, 34). Finally, we found genes with anti-apoptotic function, including several caspases, whose removal likely increases the sensitivity to apoptosis triggered by HCMV infection.

A recent study reported *PDGFRA* as the dominant hit in a CRISPR nuclease (CRISPRn) knockout screen designed to identify host factors required for HCMV entry (35), underscoring its reported role as the receptor used by HCMV strains expressing the trimeric virion glycoprotein complex (36–38). In our CRISPRi screen, *PDGFRA* knockdown conferred protection from cell death upon infection, which, however, was more pronounced using CRISPR cutting (see below, Figures 2B, S2A).

To validate and extend the host factors identified by CRISPRi, we conducted a knockout screen under the same conditions, using an established CRISPR cutting library (39) (Figure S2A, Table S2). We found good qualitative agreement of the gene-level phenotypes between CRISPRi and CRISPRn, with genes that were hits in both screening modes being either consistently protective or consistently sensitizing, but a notable variation of the phenotypes of some hits in certain pathways (Figure S2B): Hits in the knockout screen were dominated by genes involved in virus entry (*PDGFRA* and many heparan sulfate biosynthesis genes), as well as pro- and anti-apoptotic genes. In comparison, outliers in the CRISPRi screens ran a much wider gamut of biological pathways and included genes that are partially essential for host viability, independent of infection. Such genes (e.g. cullin-RING ligases and the NEDDylation machinery) were not enriched above background in the knockout screen. Our findings underscore the benefits of combining orthogonal modes of genetic screening (40). On the one hand, nonessential genes with very strong protective phenotypes (such as those involved in viral entry) seem to be more readily found in knockout screens, likely because selection pressure can act more strongly on cells with true null alleles, rather than on cells with some level of residual expression of a target gene. On the other hand, CRISPRi seems superior for resolving more weakly protective or sensitizing phenotypes, and those of essential genes. One reason may be the toxicity of DNA cutting per se, especially in a cell type with an intact p53 response (41, 42). Moreover, cells with effective knockouts of essential genes drop out of a population, leaving behind confounding cells where the intended target was not fully disrupted. In a CRISPRi screen, however, remaining cells would exhibit partial knockdown of the targets and still yield information.

## The lytic cascade resolved by single-cell transcriptomics

Our pooled screens provide a genome-scale picture of the factors involved in lytic HCMV infection, but placing them into biological pathways and linking them to a stage of the viral life cycle requires prior knowledge or dedicated follow-up experiments. To investigate the roles of critical host and viral factors systematically in more depth, we used Perturb-seq, which combines CRISPR-based genetic perturbations with a rich transcriptional readout at the single-cell level (33, 43–45). Measuring tens of thousands of single-cell transcriptomes from a population with a library of genetic perturbations provides a massively parallel way of assessing the outcome of those perturbations under uniform conditions with a high-dimensional readout. The single-cell nature of this approach additionally makes it particularly well-suited for studying viral infection, a process with great inherent variability from cell to cell, even in the absence of genetic perturbations (11, 12, 46–50).

To lay the groundwork for the Perturb-seq analysis, we first explored the natural progression of HCMV infection by recording single-cell transcriptomes from cells sampled from eight time points with two multiplicities of infection each (Figure 3A, S3A). Instead of relying primarily on synchronizing cells experimentally, which has inherent limits to its resolution due to intrinsic heterogeneity in the timing at which individual cells are infected and the rate of progression of the infection, we staged cells computationally by their transcriptional signatures. The largest sources of variability between cells were the extent of interferon signaling and the fraction of viral RNA per cell (‘viral load’), which reached levels of around 75 % (Figures S3B, C, Table S3). Cells with high fractions of viral RNA showed a marked increase in their total observed mRNA molecules (i.e. UMI counts) per cell. This indicates that the eponymous increase in cell size during infection (cyto megalo – large cell) is reflected in a higher cellular RNA content (Figures S3D, E). Together, these properties define three main subpopulations of cells: a naïve population (uninfected; IFN-negative); a bystander population (not expressing viral genes; IFN-positive); and an internally heterogeneous infected population with varying amounts of viral transcripts, which we divided into multiple subclusters (Figures 3B, S3E). Each cluster contained cells from both the low and high MOI infection samples, and the gene expression patterns between those groups of cells were extremely highly correlated (Figure S3F). This highlights both the excellent technical reproducibility of our transcriptomics workflow and that the MOI determines only the population-level response (i.e. the fraction of cells at a given stage in infection) but not the nature of the transcriptional responses in individual cells.

**Figure 3.**
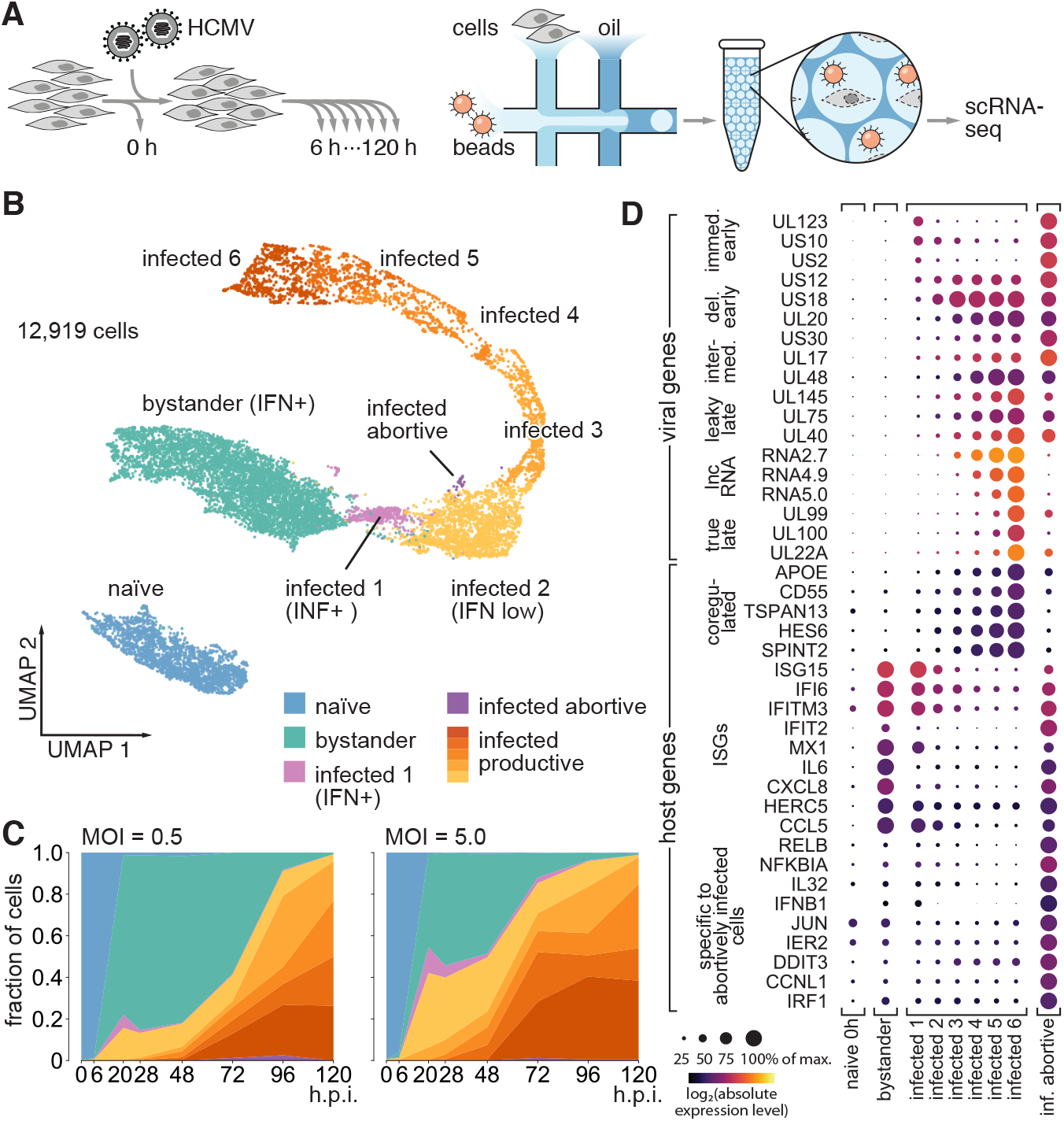
Single-cell infection time-course defines the lytic cascade of expression events as a trajectory in gene expression space. (**A**) Cells were infected with a low (0.5) or high (5.0) MOI of HCMV, harvested after times ranging from 6–120 h.p.i., pooled and subjected to emulsion-based single-cell RNA-seq (**B**) UMAP projection of the transcriptomes of all 12,919 single cells, combined from all experimental time points, color-coded by cluster membership. (**C**) Fraction of cells in each cluster as a function of time post infection and MOI. (**D**) Expression dot plot of select viral and host genes in the different clusters. Sizes of dots represent the expression normalized to the cluster with the highest expression for each gene. Colors represent absolute expression levels (scaled UMIs per gene per cell, averaged for all cells of a cluster).

Interestingly, viral gene expression and the expression of interferon-stimulated genes (ISGs) were almost entirely mutually exclusive in individual cells, a phenomenon that has been observed for HSV-1 (47). Cells with high amounts of viral loads showed entirely suppressed IFN signaling. Only cells in cluster ‘infected 1’ (Figures S3B, C, E) expressed both ISGs as well as low amounts of viral RNA, mainly classic immediate-early genes such as UL123 (IE1) (Figure 3D), indicating that these are cells in the earliest stage of infection. Together, this underscores the rapidity with which the virus effectively suppresses IFN signaling, and highlights the power of single-cell analyses in resolving this phenomenon, which may appear as concurrent expression of viral genes with ISGs in bulk measurements.

To understand the course of infection at the single-cell level, we modeled how the cell population is distributed to the different clusters as a function of time and MOI (Figure 3C, Table S3). All cells are initially in the naïve cluster. At 6 h post-infection, the first cells begin to transition to either the bystander or the infected clusters, and at 20 h.p.i. almost no naïve cells remain. The ratio of infected to bystander cells depends on the initial MOI and stays relatively constant between 20 and 48 h.p.i., with infected cells progressing to clusters with higher viral load. After 48 h.p.i, we detected another marked increase in the number of infected cells and concomitant decrease in bystander cells, corresponding to a second wave of infection, likely caused by virions released from cells that were infected early. By 96 h.p.i, even the population with low initial MOI is almost completely infected.

Among the infected clusters, the majority of cells follow a dominant trajectory with increasing viral load (clusters ‘infected 1–6’), and viral genes of known temporal kinetics show the expected patterns, e.g. true late genes are most highly expressed in the late clusters (Figure 3D, Table S3). To define the viral gene expression patterns along this trajectory at high resolution, we grouped cells from those clusters into narrow bins of 2 % of viral load and determined the profiles of all robustly quantified viral genes (Figure S4A). Many genes follow expected kinetic patterns, such as immediate-early (UL123, US10, US2) or true-late (UL99, UL100, among others). However, our high-resolution approach revealed that the (pseudo-)temporal patterns of expression of many genes were distinct from one another and did not fit the mold of the canonical ‘immediate-early’, ‘delayed-early’, ‘leaky-late’, ‘true-late’ kinetics. For instance, we identified bimodal kinetics for US6, UL78, US26, UL42 and US34, and an intermediate peak of expression for UL4 and UL48A.

Our studies reveal that the classic grouping of viral gene expression kinetics into the canonical temporal patterns does not capture the full dynamics of the HCMV viral life cycle. On the host side, a small but prominent set of host transcripts were upregulated with increasing viral load, following a pattern resembling the classic ‘leaky-late’ or ‘true-late’ kinetics (Figure 3D): *APOE, CD55, TSPAN13, HES6* and *SPINT2*. Among these, *CD55* was shown to be incorporated into budding virions to counteract the complement system (51) and its upregulation has been observed at the protein level (5). A small subpopulation, ∼ 1 % of infected cells, did not follow the dominant trajectory, but rather diverted from infected cluster 2, forming an off-trajectory where cells reach high viral loads in a distinct region of gene expression space (Figure S3B). Cells in that cluster were defined by lower UMI counts per cell, suggesting no increase in cell size (Figure S3E). Their pattern of viral gene expression was markedly different from the dominant trajectory (Figure S4B, Table S3). Immediate-early and delayed-early genes were strongly overexpressed, while true-late genes and all long noncoding RNAs were strongly depleted. As true-late gene expression depends on genome replication (52), we conclude that this trajectory is abortive. Looking at host transcripts, the abortive trajectory was characterized by lack of suppression of the interferon response. In fact, cells in the abortive trajectory not only kept responding to interferon, they were also the only cells where we detected expression of interferon-β itself (*IFNB1*), along with other cytokines and many stress response genes, prominently from the NF-κB pathway (*NFKBIA, RELB*), as well as *JUN*. Taken together, this suggests that a small subpopulation of infected cells actively secretes interferon-β and at the same time escapes suppression of the downstream response to interferon by the virus, possibly involving autocrine feedback loops in addition to paracrine signaling.

## Host-directed genetic perturbations can block or slow infection, or accelerate its progression

We next conducted a series of Perturb-seq experiments exploring the impact of targeting either host or viral factors on the viral life cycle. In contrast to the pooled screen, where phenotypes emerge by enrichment or depletion of cells over multiple days, Perturb-seq provides a high-resolution view of the impact of targeting a viral or host gene over the first 72 h post-infection, covering roughly one viral replication cycle. We first selected 52 host genes with protective or sensitizing phenotypes identified in the pooled screens, cloned them into a targeted library, along with non-targeting control sgRNAs, and delivered it into a population of fibroblasts expressing the CRISPRi machinery (Figures 4A, B, Table S4). We then challenged that population with an MOI of HCMV of 0.5 for 1 h and monitored the effects of the genetic perturbations in an average of 165 ± 50 cells per target per time point (Figure S5A). CRISPRi reduced the expression of the host targets by a median of 87 % in uninfected cells (Figure S5B) and triggered target-specific transcriptional responses (Figure S5C). In uninfected cells, we observed the strongest transcriptional responses with knockdown of IFN pathway members, LAM-TOR/Ragulator subunits, and the cullin-RING/neddylation machinery, as well as mild responses to the knockdowns of vesicle trafficking factors. The patterns of the transcriptional responses to the knockdowns organized host factors by biological pathway in a principled fashion (Figure S5C), requiring no prior knowledge and providing a layer of information that would have remained unresolved when ranking hits simply by their phenotypes in the pooled screens.

**Figure 4.**
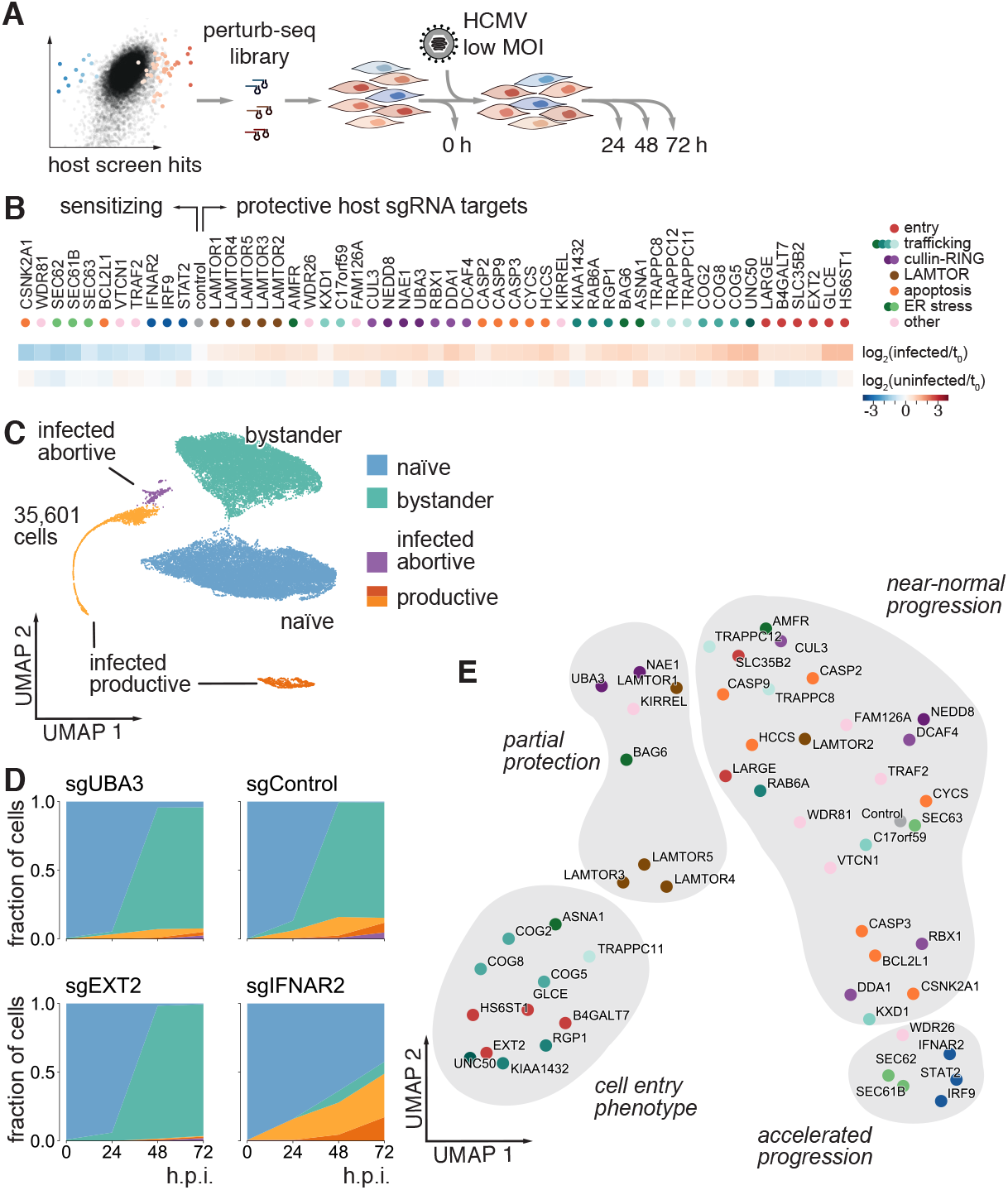
Perturbing host factors can alter the propensity of a cell to get infected. (**A**) Host dependency and restriction factors were selected from the pooled screen, cloned into a Perturb-seq library, delivered into dCas9-expressing fibroblasts, which were challenged with a MOI of 0.5 of HCMV for 24–72 h. (**B**) Selected host factors have a wide range of sensitizing to protective phenotypes, varying degrees of essentiality and cover different pathways. (**C**) UMAP projection of the transcriptomes of 35,601 cells with confidently identified sgRNAs shows the same naïve, bystander, as well as productively and abortively infected clusters found in the unperturbed infection time-course (Figure 3). (**D**) Cluster membership as a function of time post infection for cells expressing non-targeting control sgRNAs, as well as sgRNAs targeting *UBA3, EXT2* and *IFNAR2*, as representatives for the different types of responses. For a complete set of cluster membership graphs, see Figure S5H. (**E**) UMAP representation of the cluster membership data (Figure S5H) organizes host factors by their phenotypes of altered progression of infection in single cells, spanning cell entry phenotypes, partial protection from infection, near-normal progression and accelerated progression of infection.

We observed the same general split of the cell population into a naïve, a bystander, and an infected cluster, branching into a productive and an abortive trajectory (Figures 4C, S5D, E, F). The strong transcriptional response to host factor perturbations, particularly to cullin-RING/neddylation pathway members appeared as visible substructures within the naïve and bystander clusters (Figure S5G).

As expected, cells from the uninfected (0 h) sample were overwhelmingly in the naïve cluster and transitioned into the bystander and infected clusters, starting at 24 h.p.i. (Figures 4D, S5H, Table S4). The kinetics of transition of cells between the clusters was markedly different in cells with host-factor knockdowns compared to control cells. Targeting members of the heparan sulfate biosynthesis pathway such as *EXT2*, members of the COG complex, the *KIAA1432*/*RIC1-RGP1* complex, and other trafficking factors such as UNC50 efficiently prevented infection. Cells targeting the NEDD8-activating enzyme subunits *UBA3* and *NAE1* as well as LAMTOR complex members became infected, but in markedly decreased numbers. Conversely, targeting *SEC61B*, a nonessential subunit of the translocon, substantially increased the number of infected cells at the early time point of 24 h. Similarly, targeting *IFNAR2*, a subunit of the interferon receptor, as well as its downstream effectors *STAT2* and *IRF9*, increased infection rates early. Additionally, cells with perturbations of interferon pathway members failed to mount the interferon-driven transcriptional response characteristic of bystander cells and remained transcriptionally naïve as long as they were uninfected. Those cells consequently kept getting infected at much increased rates especially at later time points, when most other cells showed a robust interferon response. For a systematic classification of host targets by their progression phenotypes, we performed dimensionality reduction of the temporal cluster membership data (as shown in Figure S5H), organizing the different host factors by phenotypes on a spectrum ranging from cell entry defects to accelerated progression (Figure 4E).

Next, we sought to extend our Perturb-seq analysis to viral factors, which necessitated CRISPRn as the mode of genetic perturbation. We reasoned that when targeting a viral factor we would only obtain information from infected cells, but not from bystander or uninfected cells. We therefore challenged the cells with a high MOI of 5.0 of HCMV, without removing the inoculum, with the goal of maximizing the proportion of infected cells (Figure 5A). We selected 31 viral gene targets based on their strongly protective or sensitizing phenotypes in the pooled, virus-directed screen. Moreover, we added knockout guides targeting a representative set of 21 host factors, as well as safe-targeting guides targeting nonessential regions of the human and HCMV genomes, respectively (Figure 5B, Table S5). On average, we recovered 188 ± 77 cells per target per time point (Figure S6A). Our experimental conditions resulted in >50% of infected cells at 24 h.p.i. and a high representation of cells in the different infected subclusters (Figure 5C).

**Figure 5.**
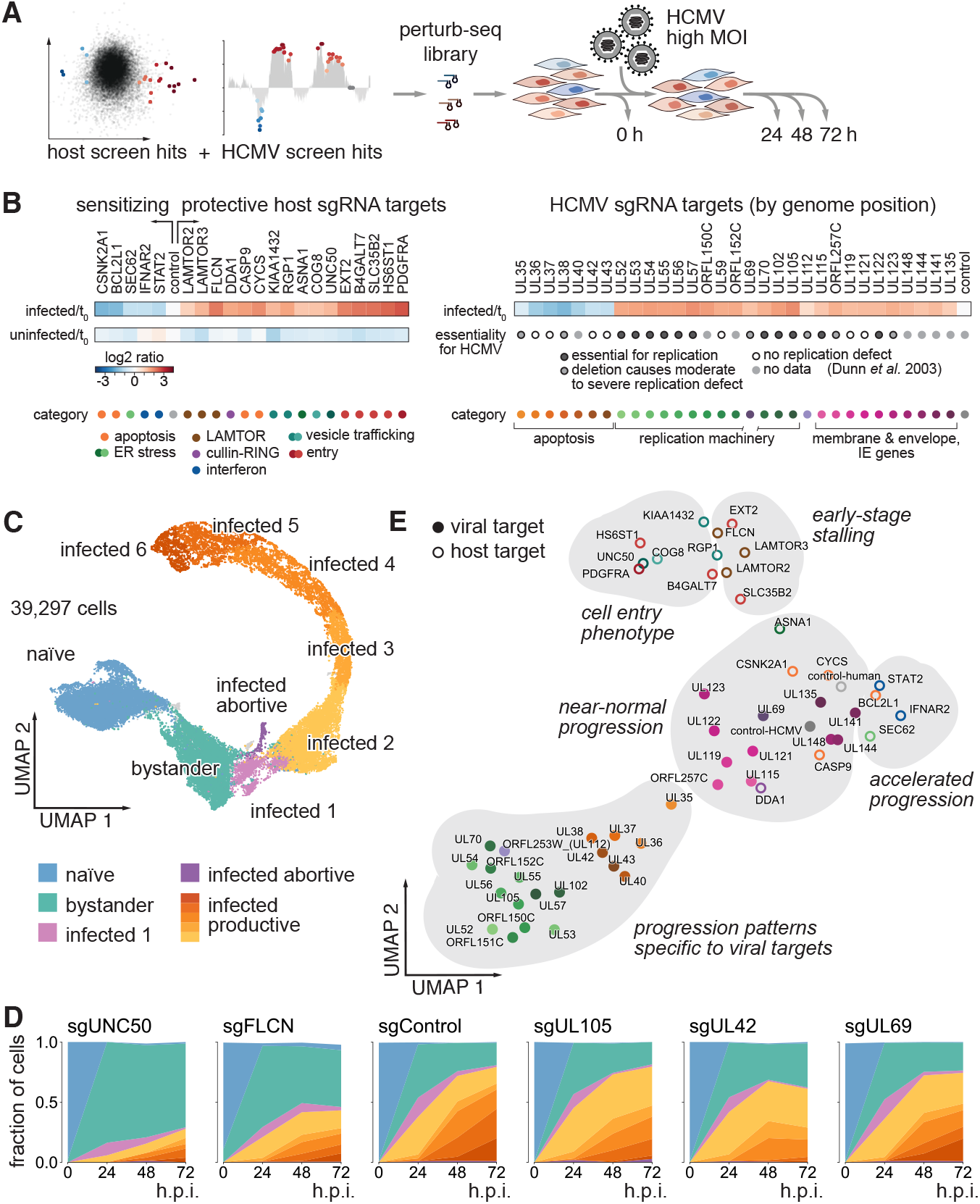
Host and virus-directed perturbations stall or accelerate progression, or shift the patterns of viral gene expression. (**A**) Host and viral factors were selected from the pooled screens, cloned into a Perturb-seq library, delivered into Cas9-expressing fibroblasts, which were challenged with an MOI of 5.0 of HCMV for 24–72 h. (**B**) Selected factors organized by their respective phenotypes in the pooled screens, essentiality for the host (determined by the uninfected arm of our pooled screen) and the virus (9), respectively, and pathway membership. (**C**) UMAP projection of the transcriptomes of 39,297 cells with confidently identified sgRNAs shows the same naive, bystander, as well as productively and abortively infected clusters found in the unperturbed infection time-course (Figure 3) and the host-directed Perturb-seq dataset (Figure 4). (**D**), Cluster membership as a function of time post infection for cells expressing sgRNAs targeting two host factors (*UNC50, FLCN*), a safe-target region of the viral genome (‘control’), as well as three viral factors (UL105, UL42, UL69), as representatives for the different types of responses. For a complete set of cluster membership graphs, see Figures S6C, D. (**E**) UMAP representation of the cluster membership data (Figures S6C, D) organizes host and viral factors by their phenotypes of altered progression of infection in single cells, spanning cell entry phenotypes, partial protection from infection, near-normal progression and accelerated progression of infection, as well as patterns specific to viral targets.

The progression of infection again varied widely depending on the targeted gene, as seen by the distributions of viral loads for each target (Figure S6B), and cluster membership of cells with a given target at the different time points (Figure 5C, S6C, D, Table S5). Host-directed knockouts confirmed our findings from the experiment where we targeted host factors by CRISPRi. The high percentages of infected cells improved the resolution of some protective phenotypes, by better distinguishing two scenarios: a reduced propensity of a cell to get infected, versus delays in progression from the early to later stages of infection (Figure S6C). For instance, knockout of *PDGFRA* (the proposed viral receptor in fibroblasts (37)) or *HS6ST1* (involved in heparan sulfate biosynthesis) almost entirely prevented infection, even in the high MOI regime. Similar levels of protection were observed in cells where *COG8* or *UNC50* were knocked out, implicating these factors as novel components required for viral entry. Conversely, perturbation of *FLCN, LAMTOR2*/*3* as well as *KIAA1432*/*RIC1* and *RGP1* permitted infection (albeit at reduced levels compared to the controls), but substantially slowed the progression of infection to late stage, indicating that these factors are essential in the early stage of infection, acting downstream of viral uncoating but prior to viral genome replication.

## Virus-directed genetic perturbations alter the trajectory of infection

Compared to targeting host genes, targeting viral genes led to qualitatively different outcomes (Figures 5D, E). Cells with virus-targeting sgRNAs generally had equal propensities to become infected (Figure S6D). This was expected because conceptually, in our experimental setup, any effect can only materialize once a cell is infected. However, cells targeting UL122 and UL123 appeared to have a slightly reduced propensity to get infected. This finding confirms the known roles of those two genes in initiating immediate-early viral gene expression, which, when suppressed, can make an infected cell present as uninfected in gene expression space.

Once infected, cells with virus-targeting sgRNAs progressed in ways specific to the target gene, evident by more complex patterns of viral load distributions (Figure S6B), and consequently of their progression through the different clusters (Figures S6D, 5D). This observation prompted us to examine in more detail the pattern of viral gene expression in infected cells, to examine whether infection follows the same trajectory in gene expression space.

On a dimensionality-reduced projection of the viral transcriptomes in infected cells, the course of infection can be visualized as a trajectory defined by a rolling average of the positions of cells with increasing viral load (Figure 6A). Cells with host-targeting sgRNAs all followed nearly identical trajectories that are congruent in shape with the default trajectory (defined by cells with control sgRNAs). However, some host-factor trajectories are necessarily shorter because some perturbations preclude cells from reaching late-stage infection (Figures 6B, C). In marked contrast, cells with virus-targeting sgRNAs occupied distinct regions in gene expression space and the corresponding trajectories diverged from the default (Figures 6D, E).

**Figure 6.**
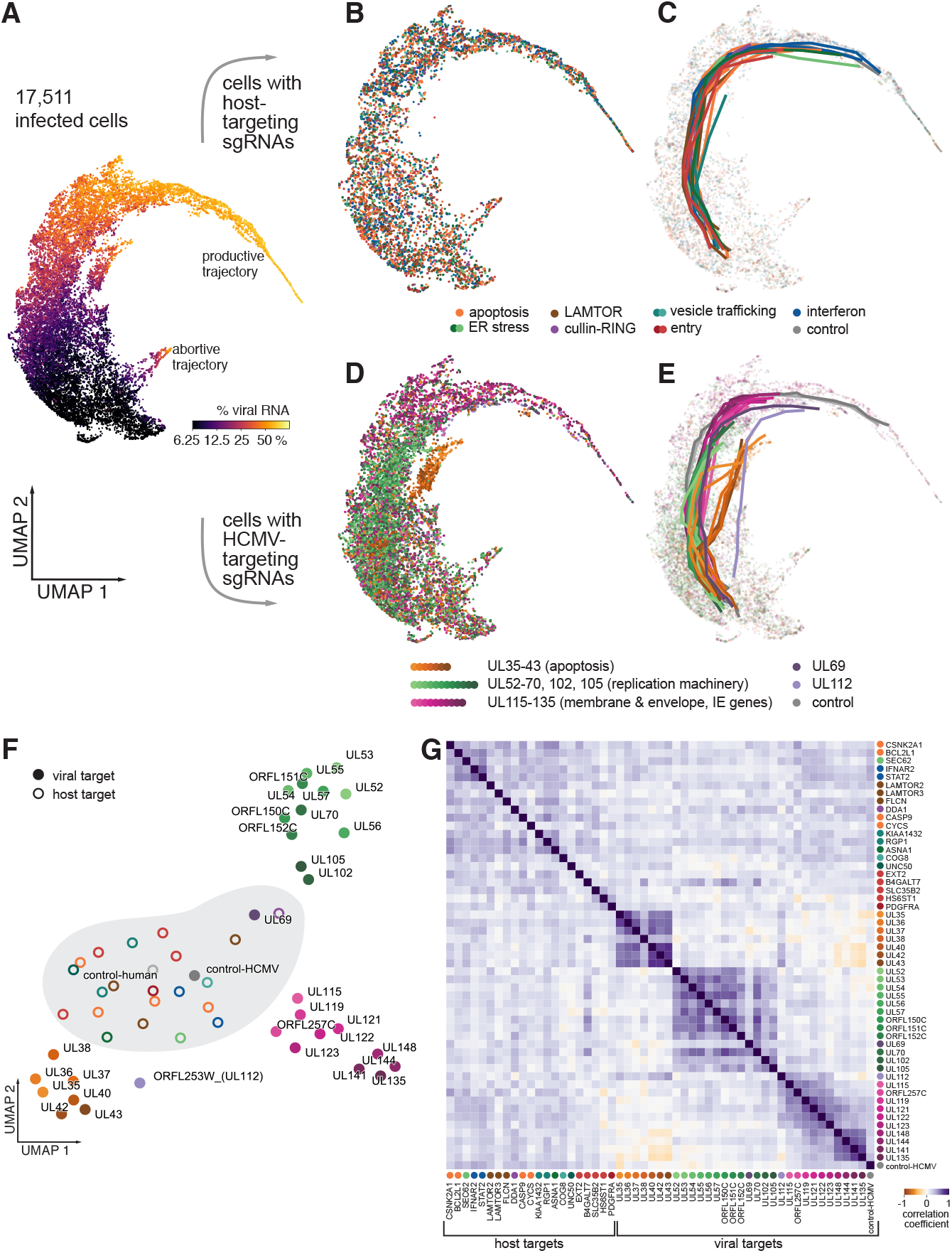
Virus-directed perturbations create alternative trajectories in viral gene expression space. (**A**) UMAP projection of the viral parts of the transcriptomes of 17,511 cells with >2.5 % viral RNA, color-coded by the fraction of viral RNA per cell. (**B**) Subsets of cells with host-directed sgRNAs, color-coded by guide identity. (**C**) Trajectories of infection for host-directed perturbations, determined by averaging the geometric position of cells with a given sgRNA target, ranked by viral load. (**D**) Subsets of cells with virus-directed sgRNAs, color-coded by guide identity. (**E**) Trajectories of infection for virus-directed perturbations. (**F**) UMAP representation of the different trajectories for each host- and virus-directed sgRNA target, calculated based on the viral gene expression matrices underlying the different virus-directed perturbation trajectories (Figure S7A) and an equivalent matrix of host-directed perturbations. All host-directed perturbations (shaded area) result in trajectories that are most similar to the control trajectories defined from both the host and virus safe-targeting controls. (**G**) Pair-wise correlation matrix of the relative viral gene expression matrices for the different trajectories highlights the three main bundles of trajectories generated by virusdirected perturbations.

To study the nature of the trajectories that infection follows in cells targeting viral factors, we quantified the expression of all viral genes along each trajectory relative to the default trajectory (defined here from cells with host-safe-targeting controls) (Figure S7A, Table S5). Cells with sgRNAs targeting non-essential regions of the viral genome followed a trajectory virtually unchanged from those with host-targeting controls, showing only mild transcriptional effects on genes in the immediate vicinity of the cut sites (within 10 kb). Reduced expression not just of the target gene itself, but also of genes located near the cut sites was a common feature for all virus-targeting sgRNAs. In addition, for all virus-targeting sgRNAs other than our safe-targeting controls, we observed widespread up- or downregulation of genes in trans, i.e. genes encoded far from the sgRNA target, indicating indirect effects on viral gene regulation caused by disruption of the target gene.

We noticed that the trajectories came in ‘bundles’ (Figure 6E), and specifically that targeting genes in the same region of the viral genome tended to result in similar patterns of deregulated viral gene expression (Figures S7A, B). This is true both for the effects in cis, which is expected, as well as for the expression changes of genes in trans, which indicates that genes are arranged in the viral genome in functional modules. To understand the relationship of the trajectories caused by targeting different genes, we projected the deviations in gene expression into two dimensions (Figure 6F) and quantified how correlated viral gene expression changes were for the different trajectories (Figure 6G). While all trajectories generated by targeting host factors were quantitatively similar to the default trajectory, viral trajectories came in three main classes: The first class of trajectories was established by targeting genes in the strongly sensitizing region (UL35–UL43). These perturbations caused reduced expression of RNA1.2 and RNA2.7, combined with overexpression to varying degrees of almost all viral genes encoded downstream of UL48, with US3 and UL54 being among the most strongly overexpressed, particularly upon targeting of UL35. Cells following these trajectories also rarely reached very high viral loads, and representation of those targets dropped substantially in the later time points (Figure S6), underscoring that these viral trajectories coincide with the cells undergoing apoptosis.

A second class corresponded to perturbations of genes in one of the strongly protective genomic modules (UL52– ORFL152C), as well as UL102 and UL105, which are located around 50 kb downstream in the genome. These genes are all involved in the viral replication machinery. Consequently, cells following these trajectories also failed to reach high viral loads, and underexpressed late viral genes, indicating a failure to replicate the viral genome upon targeting those factors.

A third class contained perturbations of genes located within the UL115–UL148 region. The corresponding viral gene expression patterns were most similar to the unperturbed trajectory outside of genes in the vicinity of the respective target genes. Of note, the kinetics of progression varied between the targeted genes within this group (Figure 5C, right panel), with perturbation of the major immediate-early transactivator genes UL122 and UL123 causing the strongest delays.

Strikingly, two viral gene targets led to trajectories of infection that were distinct from one another and from viral targets in their immediate genomic vicinity: UL69 and UL112 (ORFL253W) (Figure S6A). Both genes showed relatively weak protective phenotypes when targeted in the pooled tiling screen (Figure S1B). Further, both the UL112 and the UL69 loci encode miRNAs, which are however thought to target host and not viral genes (53). UL69 has been described as a transactivator of gene expression (54), and as involved in promoting nuclear export of unspliced RNA (55). Targeting UL69 caused a distinct viral gene expression pattern, including the downregulation of RL12, RL13, UL144, and UL155, and slight overexpression of the non-coding RNA 1.2. The UL112 locus gives rise to multiple gene products by alternative splicing, all of which we likely disrupt. Some have been shown to be involved in the recruitment of the viral polymerase activator UL44 to nuclear replication sites (56). Targeting the UL112 locus caused a viral gene expression pattern that includes the overexpression of US3, as well as upregulation of genes in the 5′ region of the genome, such as of UL22A and UL38. This pattern bears some resemblance with the pattern caused by targeting genes in the UL35–UL43 module, which sensitizes cells to cell death, while targeting UL112 is protective.

Taken together, our results show that the trajectory of infection can be derailed in defined ways by targeting viral genes, whereas targeting host factors determines the rate of progression along the default trajectory. This implies that host factors create a permissive environment, whereas viral genes orchestrate and control the viral gene expression program.

## Discussion

The waves of viral gene expression during lytic infection are a key signature of herpesvirus biology and its molecular players are subjects of intense investigation (1). Our study redefines the lytic cascade at the single-cell level as a highly resolved continuum of cellular states. We find that the large majority of cells follow this stereotypical trajectory in gene expression space, while a small but prominent subpopulation takes an alternative, abortive trajectory.

Technologically, our study breaks ground on several levels, including the use of saturating functional screens of a large viral genome, the comprehensive discovery of critical sets of genes on both sides of a host-pathogen system, and the use of single cell-analyses to understand the functional consequences of targeting these factors. Our work establishes Perturb-seq as a powerful method for functional genomics in a highly dynamic system. The single-cell approach is paramount to capturing both the inherent heterogeneity of infection in a population, and to enabling a highly multiplexed, functional analysis of genetic perturbations under defined external conditions.

Based on our high-dimensional dataset, we organized host factors both by the transcriptional responses in cells where these factors are targeted, as well as by how infection progresses in those cells. This provides a systematic classification of host factors by functional category, and revealed a range of novel factors that act in viral entry, early-stage infection, and as restriction factors. Reading out genetic perturbation phenotypes as rich transcriptional signatures also revealed that by targeting viral factors, infected cells progress along trajectories in expression space that are not normally encountered, and that do not culminate in successful replication. HCMV is entirely dependent on the transcriptional and translational machinery of its host. At the same time, our findings indicate that the lytic cascade, once set in motion, is a deterministic program that is hard-wired into the viral genome rather than reactive to the state of the cell. It will be interesting to investigate whether this dichotomy of host- versus virus-directed perturbations is a general feature of virus-host systems.

Our work provides a potential roadmap for the design of effective antiviral combination therapies by selecting sets of targets that drive the virus into distinct, nonproductive pathways while sparing or inducing apoptosis in the host, depending on the desired outcome. Similarly, our data can inform the design of engineered attenuated viral strains for vaccine development purposes. More generally, we envision that our approach of single-cell functional genomics can serve as a blueprint for studying other viruses and to define their vulnerabilities to genetic or pharmacological interventions.

## Supporting information

Table S1. HCMV tiling screen.

Table S2. CRISPRi/n host screens.

Table S3. Unperturbed single-cell time course.

Table S4. CRISPRi Perturb-seq targeting host factors.

Table S5. CRISPRn Perturb-seq targeting viral and host factors.

## ACKNOWLEDGMENTS

We thank M. A. Horlbeck for designing the HCMV tiling library, L. A. Gilbert for help setting up pooled screens, T. M. Norman, M. A. Horlbeck, J. A. Hussmann and X. Qiu for help with data analysis. A. Xu, J. A. Villalta and R. A. Pak provided technical assistance. The UCOE sequence was a gift from G. Sienski. We thank T. Fair for help with Perturb-seq experiments. We thank N. Stern-Ginossar, M. J. Shurtleff, M. Jost, R. A. Saunders, J. M. Replogle and all members of the Weissman lab for insightful discussions. J. Winkler and A. S. Puschnik provided helpful comments on the manuscript. Special thanks to O. Wueseke (impulse-science.org) for editorial help.

## Author contributions

M.Y.H. and J.S.W. conceptualized the study, interpreted the experiments and wrote the manuscript. M.Y.H. designed and carried out the experiments.

## Data and materials availability

Plasmids and libraries will be available on Addgene. Raw and processed sequencing data were uploaded to GEO (GSE165291).

## Materials and Methods

### Cell and virus culture

Human foreskin fibroblasts (HFFs; CRL-1634) and HCMV (strain Merlin; VR-1590) were purchased from the American Tissue Culture Collection. HFFs were cultured in DMEM, supplemented with 10 % FBS and penicillin-streptomycin. HCMV stocks were expanded by two rounds of propagation on HFFs and titered by serial dilution. For stable expression of the CRISPRi/n machineries in HFFs, we modified established lentiviral (d)Cas9 expression vectors (16) by inserting a minimal ubiquitous chromatin opening element (UCOE) (57) upstream of the SFFV promoter, resulting in pMH0001 (UCOE-SFFV-dCas9-BFP-KRAB) and pMH0004 (UCOE-SFFV-Cas9-BFP). The UCOE prevented epigenetic silencing that affected the original constructs.

### Pooled CRISPR screening

The HCMV tiling library was designed to contain sgRNAs targeting every single of the 33,465 PAMs in the HCMV Merlin genome (NCBI NC_006273.2), as well as 533 non-targeting controls (Table S1). It was synthesized and cloned into a lentiviral vector (Addgene #84832) as previously described (16, 17). For targeting host genes, we used the human CRISPRi v2 library (Addgene #83969) (17), and the K. Yusa et al. human knockout CRISPR v1 library (Addgene #67989) (39), respectively. Libraries were packaged into lentiviruses and delivered into (d)Cas9-expressing HFFs at an MOI of 0.3–0.5, followed by puromycin selection. Pooled screens were carried out at 500– ×1,000 coverage, i.e. 500-1,000 cells per library element per sample taken. A t_0_ sample was harvested and the remaining cells either passaged normally, or infected with HCMV at an MOI of 0.5–1.0 (for the HCMV tiling screens) or 0.1 (for the host-directed screens). Infected flasks were washed with PBS and given fresh media at days 3, 5, 7 post infection to remove dead cells, and harvested at day 7-10. Genomic DNA was extracted and digested with MfeI (pCRISPRia v2-based libraries) or HindIII (Yusa et al. library) to release a fragment containing the sgRNA cassette, followed by gel-based extraction, PCR amplification and deep sequencing as described (17). Raw count data were normalized for read depth and a small constant added to account for missing values. Phenotypes of individual sgRNAs were expressed as log2-transformed ratios of adjusted read counts between samples (Table S2). We calculated the mean of all sgRNAs specific to each host gene. For the HCMV tiling screen, we calculated a rolling average in a 250 bp window, with the average of all non-targeting sgRNAs defining the baseline.

### Single-cell RNA-seq

For the single-cell infection time course, WT HFFs were lentivirally transduced with barcoded Perturb-seq vectors to encode the experimental condition (pBA571, Addgene #85968; Table S3), followed by puromycin selection. Cells were seeded at a density of 250,000 per well of a 12-well plate and infected with an MOI of 0.5 or 5.0 with no additional media change before harvest. Infection times were staggered so that all time points for a given MOI were harvested in parallel, and pooled, aiming for roughly equal cell numbers for each time point, with a slight overrepresentation of the 20 and 28 h time points (see Figure S3D). For each MOI, pools of 10,000 cells were prepared for single-cell transcriptomics using one lane each of the Chromium Single Cell 3′ Gene Expression Solution v2 according to manufacturer’s instructions (10x Genomics), and sequenced on a NovaSeq platform (Illumina) at ∼ 100,000 reads/cell. Barcodes encoding the experimental condition were PCR-amplified from the final library and sequenced as a 5 % spike-in as described (33).

### Perturb-seq

For the host-directed CRISPRi Perturb-seq experiment, we initially selected 53 candidate genes by their strong protective or sensitizing phenotypes in the pooled screen (one gene was later removed during analysis, see below). We manually picked the two best performing sgRNAs for each candidate. Additionally, we added six control constructs targeting GFP (which is not present in our HFFs). For the host- and virus-directed CRISPRn Perturb-seq experiment, we selected a set of 21 host factors, of which 19 were already among the targets CRISPRi Perturb-seq experiment, had no strong essentiality knockout phenotypes and comparable protective or sensitizing phenotypes in both the pooled host-directed CRISPRi and CRISPRn screens (see Figure S2). We further added *PDGFRA* and *FLCN*, both of which were strong hits in the pooled CRISPRn screen. For each host target, we manually picked the two best performing sgRNAs from the pooled screen. In addition, we selected 31 viral targets with strong protective or sensitizing phenotypes, corresponding to the three strongest modules identified in the HCMV tiling screen (see Figures 1B, S1). From the tiling screen, we selected the two highest-ranking sgRNAs for each target gene based on the following scoring system: From the pool of unique sgRNAs falling within the gene boundaries and having a Doench score (58) of >0.5, we calculated the absolute average phenotype across replicates and subtracted a penalty defined as the difference between replicates plus the average absolute essentiality phenotypes on a log2 scale. We designed a number of safe-targeting control sgRNAs targeting intergenic DNA in the US2-US12 region. This region was selected based on its near-neutral phenotypes in the tiling screen (Figure S1), a lack of essential genes (8, 9), and its comparatively large spaces between consensus genes. Further, in some bacterial artificial chromosome (BAC) constructs harboring HCMV genomes, this region has been replaced by the BAC backbone, underlining its non-essential nature during infection in tissue culture (59). We picked five sgRNAs based on their Doench scores from a pool of unique sgRNAs targeting the intergenic regions and having survival and essentiality phenotypes of <0.5 (log2 scale) in all replicates. In addition, we included four control sgRNAs directed against safe-harbor loci in the host genome, which we repurposed from gene knock-in applications.

All sgRNAs were synthesized as individual oligo pairs (IDT) and cloned into a barcode library containing plasmid pool (pBA571, Addgene #85968), thereby linking each sgRNA to a unique guide barcode contained within the 3′-UTR of the puromycin resistance gene (33). Barcodes were validated to not contain homo-oligomers or sequences resembling transcription termination signals. All sgRNA and barcode sequences are listed in Tables S4 and S5. sgRNAs vectors were individually packaged into lentiviruses, titered separately and pooled to ensure equal representation. This workflow prevents scrambling of guide sequences and associated barcodes by recombination, which is a concern in pooled lentivirus preparations (60). We delivered the pooled library into (d)Cas9-expressing HFFs at an MOI of 0.3 followed by puromycin selection. Cells were seeded at 250,000 per well of a 12-well plate and infected with HCMV at an MOI of 0.5 for 1 h, followed by media exchange (for the CRISPRi host-directed experiment) or an MOI of 5.0, leaving the inoculum on the cells, with the goal of maximizing the numbers of infected cells (for the CRISPRn host and virus-directed experiment). Cells were harvested in the uninfected state (designated as 0 h) and at 24, 48 and 72 h.p.i. We aimed at a representation of each library element by around 100 cells per time point (for actual cell numbers see Figures S4A, S5). Cells were collected and prepared for sgRNA-seq using the 10x Chromium platform as described above for the single-cell infection time course. Libraries were sequenced on a HiSeq4000 (Illumina) at ∼40,000 reads/cell.

### Single-cell data analysis

Cells were collected and prepared for sgRNA-seq using the 10x Chromium platform as described above for the single-cell infection time course. Libraries were sequenced on a HiSeq4000 (Illumina) at 40,000 reads/cell. Single-cell data analysis Raw sequencing data were submitted to ‘cellranger’ v2.0.1 (10x Genomics) according to the manufacturer’s instructions. We compiled a reference transcriptome from the hg19 human genome and a custom assembly of HCMV coding transcripts based on our previous ribosome profiling dataset (4) as distributed as part of the ‘Plastid’ python library demo dataset (61). We manually added four well-established lncRNA transcripts (RNA1.2, 2.7, 4.9, 5.0). Internal ORFs were removed as they would create ambiguous mappings, as were ORFs overlapping with the aforementioned lncRNAs. Cells retained in the final dataset had to cross the default cellranger quality thresholds, as well as have one unique lentiviral barcode assigned with high confidence (33). During data analysis of the perturb-seq experiments, three CRISPRn sgRNAs targeting host genes were removed computationally because they were found to be inactive, as seen by lack of transcriptional responses and viral load patterns similar to cells with control sgRNAs. One host gene, *RBBP5*, was similarly excluded from both the CRISPRi and CRISPRn datasets as it became apparent that its knockdown/knockout causes differentiation of cells and a strong transcriptional response, rather than true protection against infection (see Tables S4 and S5).

Percentages of viral RNA (viral loads) were calculated as the fraction of total UMIs per cell mapping to viral genes.

Gene expression was normalized in each cell by a factor scaling the total UMIs mapping to human transcripts to its average number across all cells in a population. This accounts for the fact that infected cells have much higher total UMI counts, indicating that viral transcripts go ‘on top’ of human transcripts (see Figure S3E).From the unperturbed time-course experiment, we defined a set of robustly detected genes as those with >10,000 UMIs total across all cells in that population (3,588 genes in total, of which 106 are viral genes). This set of expressed genes was used consistently for the analysis of all whole-transcriptome single cell datasets in this study. For heatmap representations of gene expression as a function of viral load, cells were binned by viral load and gene-level expression values averaged in each bin. Bin widths of 2 % or 10 % were selected depending on the available number of cells. We visualized a slightly larger set of viral genes, namely those expressed in >95 % of cells in at least one of these 2 % viral load bins (114 genes in total). Viral transcriptome-centric trajectory analyses (Figures 6, S7) were also based on this set of viral genes.

Cell cycle phases were scored based on marker genes as described (33). Using a similar approach, we calculated an IFN score by summing (in each cell) and subsequently z-scoring (across cells) the normalized expression values of the following set of robustly quantified ISGs (*PSMB8, PSMB9, PSME1, PSME2, ISG15, ISG20, IRF7, MX1, MX2, GBP1, GBP2, GBP3, IFI6, IFI44, IFI35, IFI16, IFI27, IFIH1, IFI44L, IFIT1, IFIT2, IFIT3, IFIT5, IFITM1, IFITM2, IFITM3, EIF2AK2, OAS1, OAS2, OAS3, CNP, PLSCR1, BST2, BTN3A2, XAF1, CASP1, CASP4, CASP7, GSDMD*).

To visualize single-cell datasets, we performed dimensionality reduction by UMAP (62) based on the matrix of scaled expression values of the set of robustly detected genes (host + viral genes in Figures 3, 4, S3, S4; viral genes in Figures 6, S6). Clusters of cells were defined by Leiden clustering (63) or HDBSCAN (64). To determine trajectories, selected cells were ranked by viral load and the geometric position of cells averaged in a sliding window that was shifted in increments of 0.2 window sizes. Window sizes were selected based on the total number of available cells: 100 cells for each sgRNA targets; 500 cells for cells with control sgRNAs. UMAP was also used for a dimensionality-reduced visualization of the similarities of the cluster membership data as a function of time and sgRNA target (Figures 4, 5E, underlying data in Tables S4, S5) and of the viral gene expression data along the trajectories defined by cells with individual sgRNA targets (Figure 6F, underlying data in Table S5).

## Supplemental Tables

**Table S1. HCMV tiling screen**.

- sgRNA sequences of the HCMV tiling library; raw sequencing counts in the screen, and normalized guide-level phenotypes.
- Gene-level phenotypes for consensus genes (based on NCBI).
- Gene-level phenotypes for all ORFs, based on (4).

**Table S2. CRISPRi/n host screens**.

- Raw sequencing counts for the human genome-wide CRISPRi screens.
- Gene-level phenotypes for the human genome-wide CRISPRi screens.
- Raw sequencing counts for the human genome-wide CRISPRn screen.
- Gene-level phenotypes for the human genome-wide CRISPRn screen.

**Table S3. Unperturbed single-cell time course**.

- Metadata annotations for all cells in the final dataset.
- Table of the expressed barcodes used to deconconvolve the pooled cells into the experimental time points.
- Expression values of all robustly detected host and viral genes in the individual clusters.
- Expression values of robustly detected viral genes along the default trajectory of infection.
- Expression values of viral genes along the abortive trajectory of infection.

**Table S4. CRISPRi Perturb-seq targeting host factors**.

- Metadata annotations for all cells in the final dataset.
- sgRNA sequences, guide barcodes, and annotations for all elements of the library.
- Table of cell numbers in each cluster as a function of experimental time.
- Expression values of all robustly detected host genes in the naïve cluster, as a function of sgRNA target.
- Expression values of all robustly detected host genes in the bystander cluster, as a function of sgRNA target.

**Table S5. CRISPRn Perturb-seq targeting viral and host factors**.

- Metadata annotations for all cells in the final dataset.
- sgRNA sequences, guide barcodes, and annotations for all elements of the library.
- Table of cell numbers in each cluster as a function of experimental time.
- Expression values of all robustly detected viral genes along the trajectories of infection as a function of sgRNA target.

**Figure S1. related to Figure 1.**
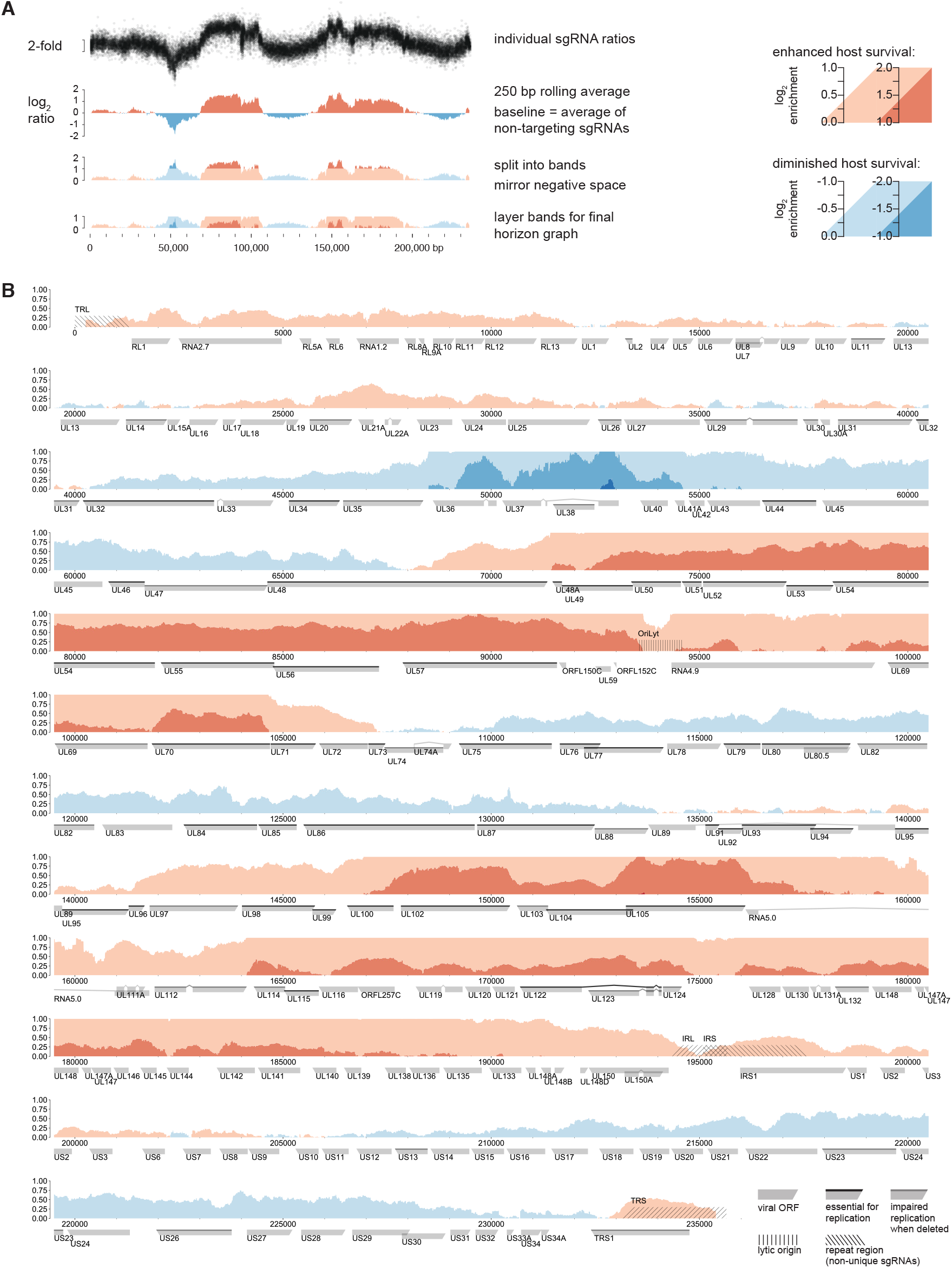
**(A)** Data processing for the HCMV tiling screen. We calculated log2 ratios of each individual sgRNA in the surviving over the t_0_ populations. Ratios were averaged in a sliding 250 bp window. The average of the ratios of the non-targeting sgRNA population was set as the baseline. The plot was then colored based on the sign of the average phenotype and layered in bands of decreasing lightness, one log2 unit wide. The negative space was mirrored on the baseline, and bands were stacked for the final horizon plot representation (65). **(B)** High-resolution Horizon graph of the phenotypic landscape of the HCMV genome. Shades of blue denote sensitization to host cell death, shades of red denote protection from cell death upon HCMV genome cleavage. Major features of the HCMV genome are annotated. sgRNAs targeting internal and terminal repeat regions (hashed) typically have multiple target sites and likely result in higher-order fragmentation of the HCMV genome, exacerbating their respective phenotypes. Viral ORFs are classified by their essentiality for viral replication based on (9). ORFL150C, ORFL151C (originally named UL59, but thought to not be expressed as a protein (66), causing it to be dropped from the consensus annotation), and ORFL152C were the only short ORFs with strong phenotypes in areas of the genome devoid of consensus genes. UL48 was the only gene that showed a substantial phenotype gradient within its gene body: Cutting the N-terminal region caused mild sensitization to death upon infection, whereas cutting the C-terminus had the opposite effect.

**Figure S2. related to Figure 2.**
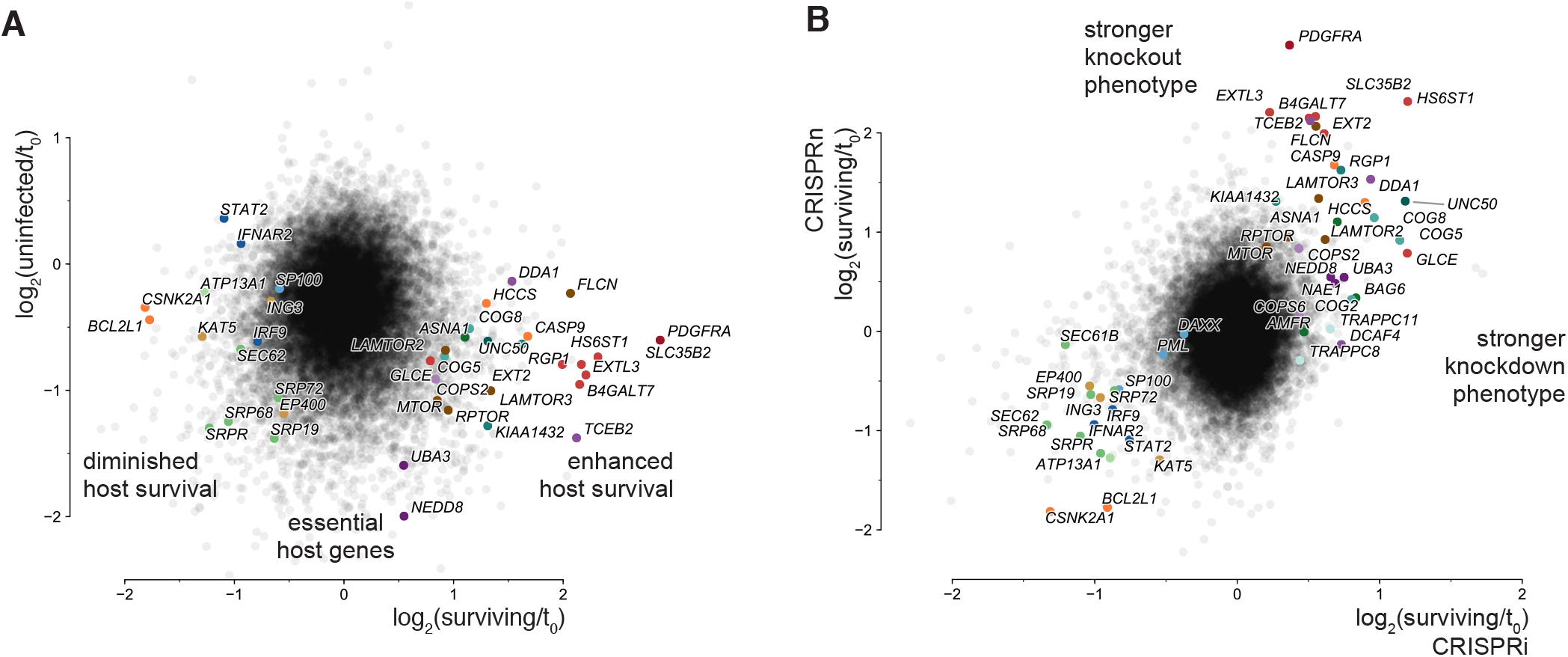
(**A**) Results of host-directed CRISPRn screen displayed as a scatter plot of average gene essentiality (i.e. infection-independent phenotype; y-axis) vs. protection/sensitization to death upon HCMV infection (i.e. infection-dependent phenotypes; x-axis). Note that due to the experimental design of the screen, the apparent gene essentiality phenotypes are underestimating the real essentiality because t_0_ refers to the beginning of HCMV infection, not lentiviral delivery of the sgRNA library. (**B**) Direct comparison of CRISPRi and CRISPRn phenotypes for host targets represented in both libraries. Hits involved in viral adhesion and entry, as well as host cell survival or apoptosis are more pronounced in the CRISPRn screen. Cullin/RING pathway members and some vesicle trafficking factors were only resolved in the CRISPRi screen.

**Figure S3. related to Figure 3.**
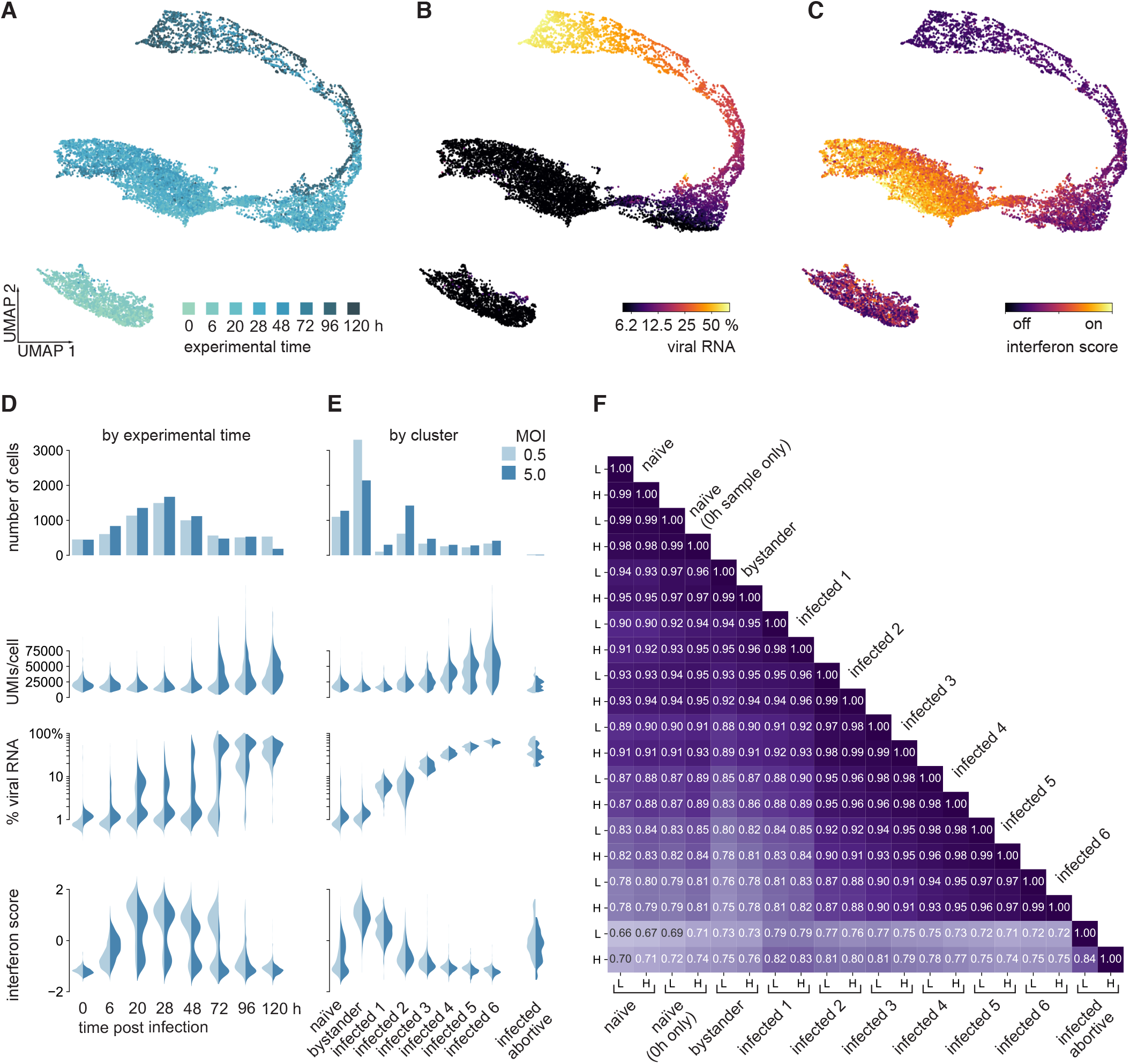
(**A, B, C**) UMAP projections of single-cell transcriptomes (same as in Fig. 3B), color-coded by experimental time post infection (A), percentage of viral transcripts per cell (B) and interferon score, calculated from the normalized expression of interferon stimulated genes (C). (**D, E**) Numbers of cells, as well as distributions of UMIs per cell, percentage of viral transcripts per cell, and interferon score, broken down by cells for each MOI and each experimental time point (D), and for cells for each MOI and cluster membership (E). (**F**) Pearson’s correlation matrix of gene expression values (average logarithmized, scaled UMIs per gene per cell) for all clusters, broken down by low (L) and high (H) MOI conditions.

**Figure S4. related to Figure 3.**
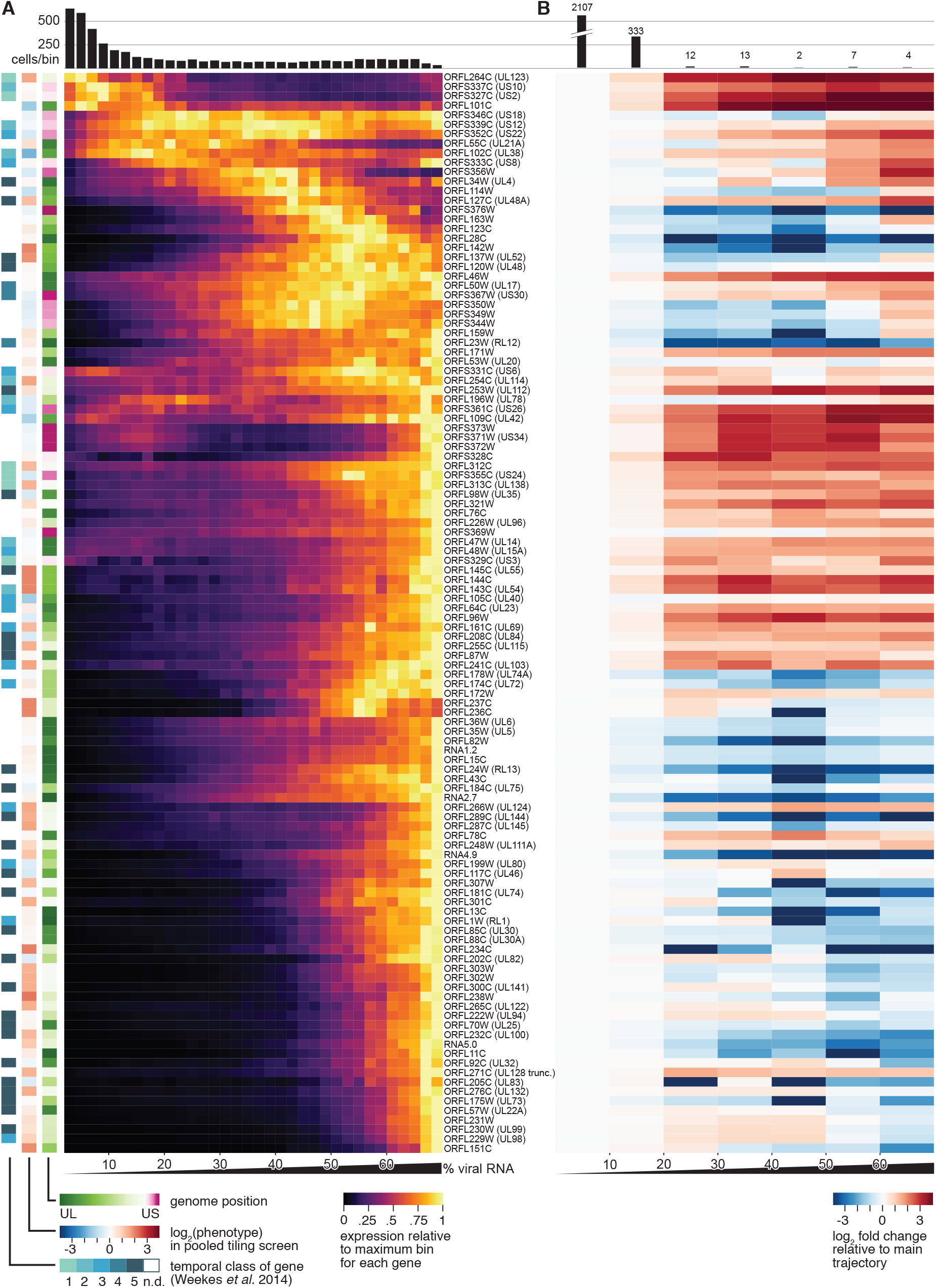
(**A**) Gene expression profiles for robustly detected viral genes along the dominant trajectory (clusters ‘infected 1-6’). Cells were grouped in bins spanning 2 % of viral RNA and the gene expression (scaled UMIs per gene per cell) averaged for all cells in each bin. The heatmap shows the expression relative to the highest bin. Individual viral genes are ordered by similarity of the profiles, and annotated by genome position, phenotype of cutting within the body of a gene in the pooled virus-directed CRISPR screen (see Figs. 1 and S1), and by the temporal profile as determined in a bulk proteomics study (5). Note the relationship between a gene’s temporal class and its phenotype in the pooled screen: True-late and leaky-late genes predominantly showed protective phenotypes, whereas earlier classes also contained sensitizing genes. (**B**) Gene expression profiles of viral genes along the abortive trajectory (clusters ‘infected 1–2’ and ‘infected abortive’). Cells were grouped in bins spanning 10 % of viral RNA and the gene expression averaged for all cells in each bin. The heatmap shows the expression relative to the expression in an equivalent bin of the dominant trajectory.

**Figure S5. related to Figure 4.**
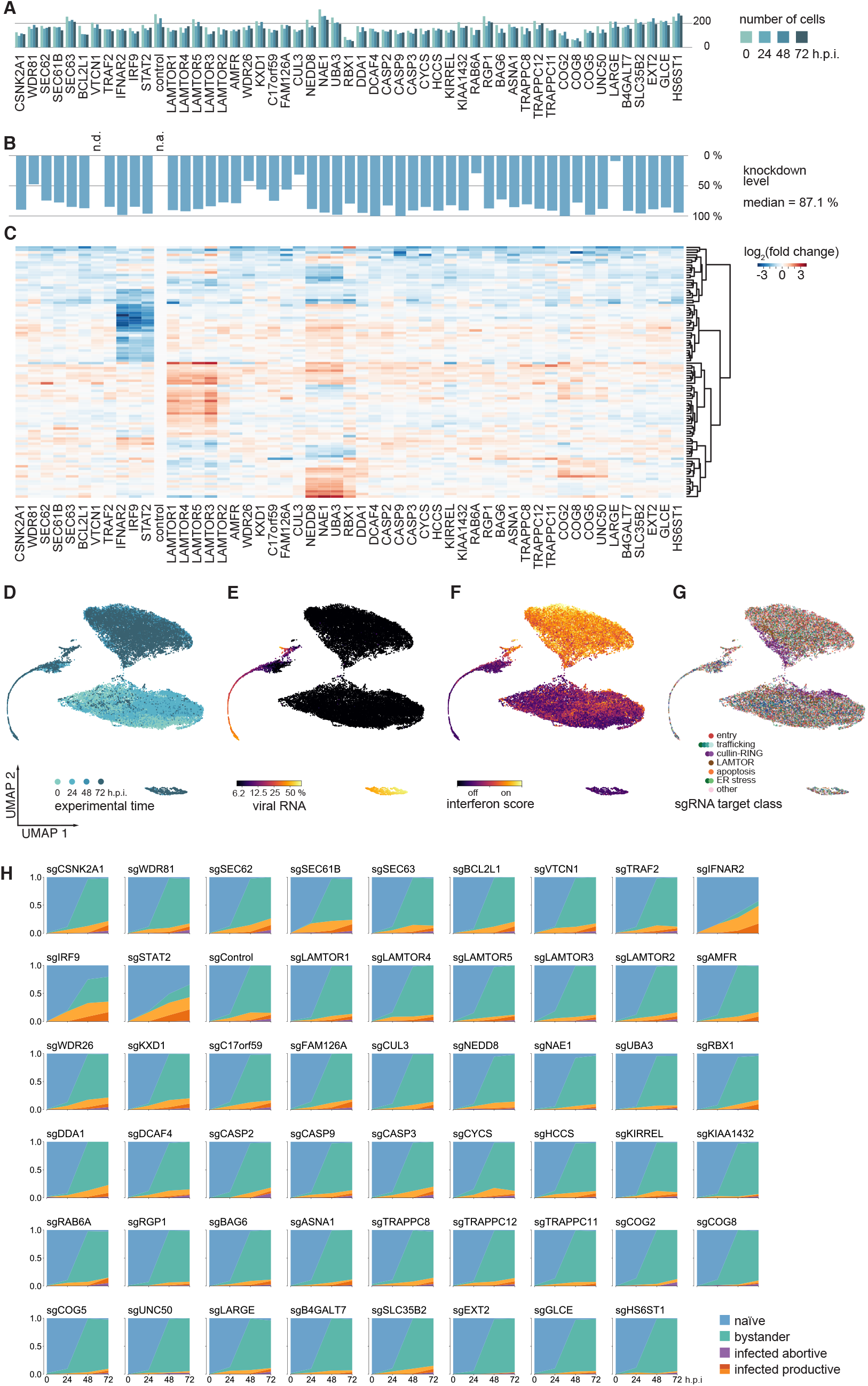
(**A**) Numbers of single cells for each sgRNA target for each experimental time point in the host-directed CRISPRi Perturb-seq experiment. The average is 165 ± 50 (mean ± standard deviation) cells per sgRNA per time point. (**B**) Knockdown levels for each sgRNA target calculated from the expression of the target gene in cells with a given sgRNA target relative to cells with control sgRNAs. No transcript at all was detected for VTCN1. Median knockdown level was 87.1 %. (**C**) Hierarchical clustering of expression changes of the most variable 100 genes (excluding the targeted factors) in response to host factor knockdown in naïve cells, relative to naïve cells with control sgRNAs. (**D, E, F, G**) UMAP projections of single-cell transcriptomes of cells from the host-directed Perturb-seq experiment (same as in Fig. 4C), color-coded by experimental time post infection (D), percentage of viral transcripts per cell (E), interferon score, calculated from the normalized expression of interferon stimulated genes (F), and by pathway of the targeted host factor in each cell (G). (**H**) Cluster membership as a function of sgRNA target and time post infection.

**Figure S6. related to Figure 5.**
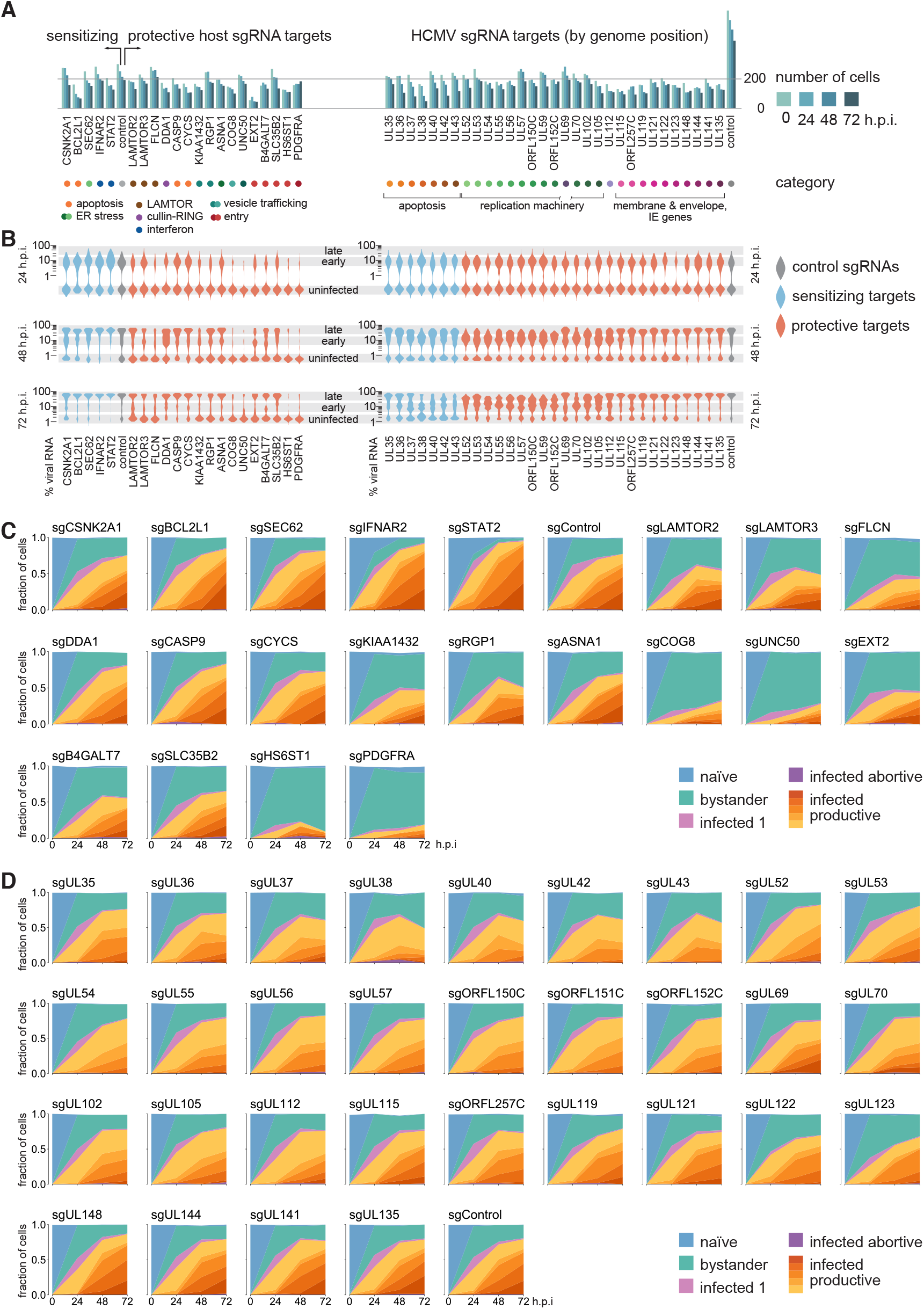
(**A**) Numbers of single cells for each sgRNA target for each experimental time point in the host-directed CRISPRi Perturb-seq experiment. The average is 188 ± 77 (mean ± standard deviation) cells per sgRNA per time point. Note the over-proportional drop in numbers in late time points of cells with apoptosis-related sgRNA targets. ‘Control’ denotes all safe-targeting sgRNAs, which are 4 and 5 distinct sgRNAs targeting the host and virus, respectively. (**B**) Violin plots of the distribution of viral RNA fraction per cell as a function of time post infection and the sgRNA target (red, protective phenotype; blue, sensitizing phenotype; grey, control). Regions of the violin plot corresponding to uninfected cells, as well as early and late stages of infection are highlighted. Note that uninfected cells have non-zero background amounts of viral RNA, and those background levels are higher in later time points, indicating leaking of viral RNA from dying cells. (**C, D**) Cluster membership as a function of sgRNA target and time post infection for cells with host-targeting sgRNAS (C) and virus-targeting sgRNAs (D).

**Figure S7. related to Figure 6.**
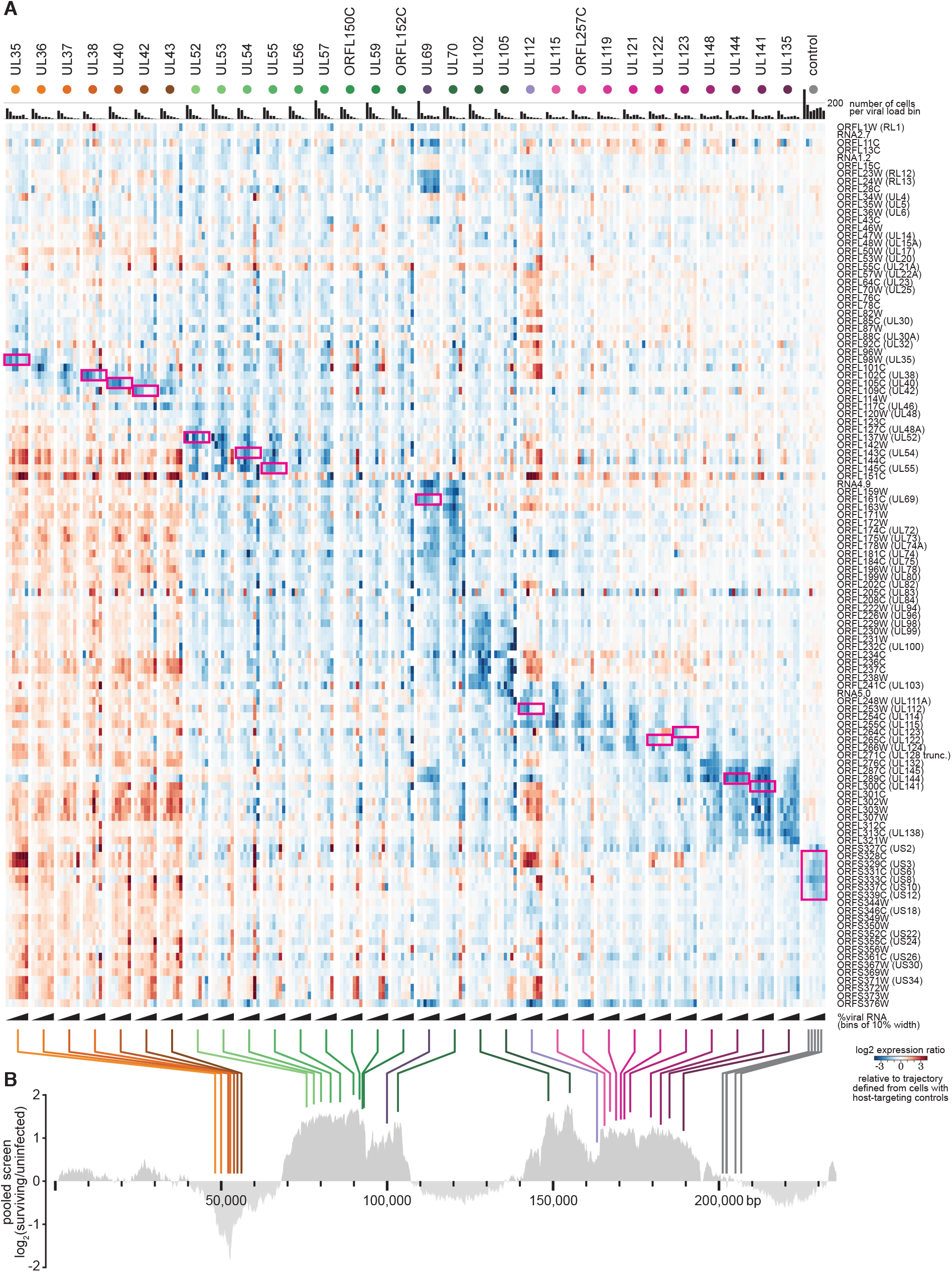
(**A**) Heatmaps of viral gene expression for all cells with virus-targeting sgRNAs. For each sgRNA target, cells were grouped in bins of 10 % of viral RNA fraction, and the expression of viral genes plotted relative to a corresponding bin defined by cells with host-directed, safe-targeting sgRNAs (similar to Figure S3H), representing the unperturbed, dominant trajectory. Both the columns (viral sgRNA targets) as well as the rows (expressed viral genes) are ordered by genome position. This facilitates the distinction of gene expression effects in cis, i.e. the immediate effect of cutting on genes adjacent to the cut site, as opposed to in trans, which are reflecting an altered trajectory of infection. Pink boxes indicate the sgRNA target genes. (**B**) Mapping the sgRNA targets onto the phenotypic landscape of the HCMV genome (see Figure 1B), indicating genome position and phenotype in the CRISPRn tiling screen.

